# Label-Free Density-Based Detection of Adipocytes of Bone Marrow Origin Using Magnetic Levitation

**DOI:** 10.1101/462002

**Authors:** Oyku Sarigil, Muge Anil-Inevi, Gulistan Mese, H. Cumhur Tekin, Engin Ozcivici

**Affiliations:** Department of Bioengineering, Izmir Institute of Technology, Urla, Izmir, Turkey 35430; Department of Molecular Biology and Genetics, Izmir Institute of Technology, Urla, Izmir, Turkey 35430

**Keywords:** magnetic levitation, bone marrow stem cells, cell detection, adipocytes, single cell density measurement

## Abstract

Adipocyte hypertrophy and hyperplasia are important parameters in describing abnormalities in adipogenesis that are concomitant to diseases such as obesity, diabetes, anorexia nervosa and osteoporosis. Therefore, technical developments in detection of adipocytes become an important driving factor in adipogenesis research. Current techniques such as optical microscopy and flow cytometry are available in detection and examination of adipocytes, driving cell- and molecular-based research of adipogenesis. Even though microscopy techniques are common and straightforward, they are restricted for manipulation and separation of the cells. Flow cytometry is an alternative, but mature adipocytes are fragile and cannot withstand the flow process. Other separation methods usually require labeling of the cells or usage of microfluidic platforms that utilize fluids with different densities. Magnetic levitation is a novel label-free technology with the principle of movement of cells towards the lower magnetic field in a paramagnetic medium depending on their individual densities. In this study, we used magnetic levitation device for density-based single cell detection of differentiated adipogenic cells in heterogeneous populations. Results showed that magnetic levitation platform was sensitive to changes in lipid content of mesenchymal stem cells committed to adipogenesis and it could be successfully used to detect adipogenic differentiation of cells.

## INTRODUCTION

Adipose tissue, once considered only as a mechanical support and insulating tissue serving mainly for energy storage, is now known to support crucial endocrine and immune functions in various tissues ^1-4^. For example, abnormal adipogenesis in bone marrow can be observed in several diseases such as obesity, diabetes, anorexia nervosa and osteoporosis ^5-11^. Marrow adipocytes are derived from mesenchymal stem cells (MSCs), which also have the capability of differentiation into osteoblasts ^12-14^. Once a marrow adipocyte is formed, it can interfere with cellular differentiation, bone remodeling and hematopoiesis via secreted factors ^6, 11, 15, 16^ as there is a negative relationship between bone marrow adipocytes and bone density ^17, 18^. Disturbed balance of adipogenesis and osteoblastogenesis in the bone marrow influences bone formation, and increased marrow adipocytes inhibit bone reformation ^19-24^. Moreover, bone fragility increases with skeletal aging that is characterized by a shift of balance between adipogenesis and osteoblastogenesis to increased adipogenesis ^17, 25, 26^. Marrow adipogenesis can further be linked to changes in immune and endocrine system functions ^27-30^.

Complex functions of adipocytes with their strong association to the epidemiology of obesity and obesity-related diseases have remarkably increased the interest in morphology and physiology of adipocytes ^31^. In this context, it has become critical to detect and identify adipocytes to for hypertrophy (increasing cell size), hyperplasia (increasing cell number) and thus to examine the underlying mechanisms of the formation and development of adipose tissue in healthy and abnormal tissues ^32, 33^. Adipocyte detection on cellular level relies on the formation of intracellular lipid droplets and an increase in cell volume ^34-36^. Optical microscopy is commonly used to visualize adipocyte differentiation by staining the cells with lipophilic dyes such as Oil Red O, Nile Red and Sudan Red ^37-39^. Although optical microscopy is common and useful for detection and examination of the monolayer lipid-accumulated cells, produced results mainly lead to a qualitative assessment. Adipogenic differentiation can also be observed through molecular markers of adipogenesis such as P107, PPARγ, C/EBPα and aP2 ^23, 40-42^, however, these techniques are time-consuming, labor-intensive and do not allow the recovery of cells for further studies. Advanced techniques, such as flow cytometry, can be used to detect differentiated adipocytes ^35^. However, nozzles in these systems limit the application to pre-adipocytes, as mature adipocytes are fragile and cannot withstand the flow ^43-45^. Alternative cell separation platforms are available based on fluorescent or magnetic markers ^46-48^, or utilization of complex microfluidic platforms that either use fluid interfaces where cells are stratified on different fluids based on their densities ^49^. Furthermore, sensors that can track changes in electric cell-substrate or using Coherent anti-Stokes Raman scattering (CARS) imaging can be used as label-free methods ^50-52^, however, these techniques are relatively expensive, require complex instrumentation and cannot perform single cell detection.

Cell density can be utilized as an indicator of cell state such as apoptosis, disease state, cell cycle ^53-56^. Changes in cell density are also observed during cellular differentiation ^57^. The conventional method to determine cell density is via density gradient centrifugation ^58, 59^. Even though this technique is relatively simple and cost-effective, it may cause cell damage due to high centrifugal forces, can only yield the average density of target cell population and cannot discriminate small differences in cell densities ^49, 60^. Density measurements based on a single cell are possible through microfluidic technology, such as suspended microchannel resonator (SMR) system ^57, 58^ and optically induced electrokinetics (OEK) microfluidic platform ^61^. Although SMR system is very precise in measuring single cell density, it is time-consuming and compatible liquids with different densities need to be selected specifically to the application ^62^. OEK platform, on the other hand, has a complex design with optical elements and electrode. An alternative methodology for single cell density measurements is the magnetic levitation technology that allows real-time, label-free separation of cell populations with the principle of movement of cells towards the lower magnetic field in a paramagnetic medium based on their density ^60, 63, 64^. Magnetic levitation system has been previously applied with different shape and configurations of magnets to improve accuracy and throughput in detection of object/cell densities ^65-67^.

In this study, a quantitative method was demonstrated to detect the changes in the single cell density of lipid-accumulated bone marrow cells during adipogenic differentiation by using magnetic levitation principle. We showed that magnetic levitation platform was sensitive to changes in lipid content of mesenchymal stem cells committing to adipogenesis and it could be successfully used to detect adipogenic differentiation of cells. We believe this quantitative, cost-effective and label-free microfluidic system serves as a potential method to be applied in future adipocyte and lipid research.

## RESULTS

### Calibration of the magnetic levitation system with polymeric beads

In this study we used a custom designed microfluidic platform that is composed of a holder, two opposing neodymium magnets, a capillary channel that can hold cells in a paramagnetic medium, and two parallel mirrors to visualize cell levitation (Figure 1A). The platform can generate magnetic forces (F_mag_) that can balance the corrected gravitational force (F_g_) which is the combination of gravity and buoyancy force vectors (Figure 1B).

**Figure 1.**
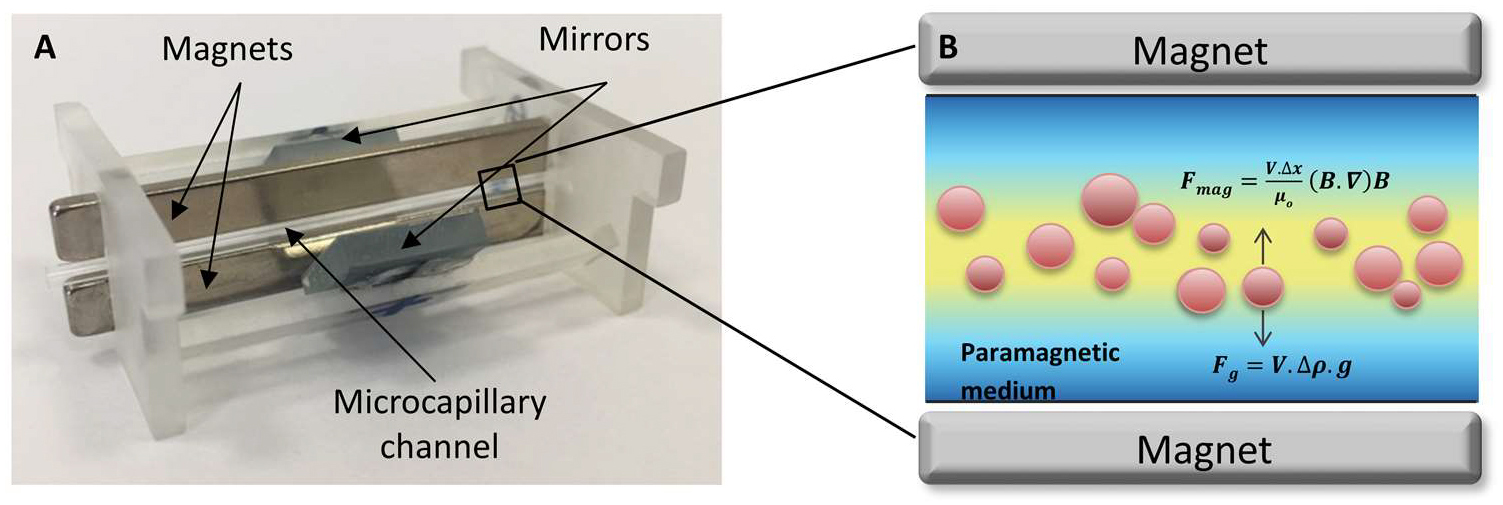
Magnetic levitation setup and working principle. (A) Structure of magnetic levitation device composed of two neodymium magnets, microcapillary channel and two mirrors placed at 45°. (B) Forces acting on cells at equilibrium position in the device; where F_mag_: magnetic force, F_g_: corrected gravitational force, V: cell volume, Δx: magnetic susceptibility difference between paramagnetic medium and cell, μ_0_: magnetic permeability of free space, B: magnetic field, Δρ: density difference between paramagnetic medium and cell, g: gravity force.

Prior to levitation of cells, microfluidic device was calibrated for density-based detection of adipogenesis via determining the levitation heights of polymeric beads with known densities (1 g/mL, 1.02 g/mL and 1.09 g/mL) in the culture medium containing paramagnetic Gadolinium (Gd^3+^) at 25 mM, 50 mM and 100 mM concentrations (Figure 2A). Increased bead density and decreased Gd^3+^ concentrations lead to lower levitation heights, which was measured as average bead distance from the top surface of the bottom magnet. Levitation heights of beads with 1.09 g/mL were 37.15%, 22.86% and 13.76% lower than the beads with 1 g/mL in the medium containing 25 mM, 50 mM and 100 mM Gd^3+^ (all p<0.05), respectively (Supplementary Figure S1). Furthermore, increased Gd^3+^ concentration decreased the difference of levitation height between different densities of polymeric beads defining the working span of the microfluidic device. The difference between the levitation heights of beads (range) with densities of 1.09 g/mL and 1 g/mL was found as 425 μm, 261 μm and to 156 μm (Figure 2A; blue arrows), when levitation was performed with 25 mM, 50 mM and 100 mM Gd^3+^, respectively. Similarly, increased Gd^3+^ concentration decreased the range between beads with densities of 1.09 g/mL and 1.02 g/mL from 346 μm to 203 μm and to 116 μm (Figure 2A; red arrows). Inverse relation of levitation height and bead density values showed a strong correlation (R^2^ > 0.99) for different Gd^3+^ concentrations (Figure 2B) with the linear fits ρ = - 0.000209001 × h + 1.240573 for 25 mM Gd^3+^; ρ = -0.000344456 × h + 1.393534 for 50 mM Gd^3+^; and ρ = -0.000583928 × h + 1.659352 for 100 mM Gd^3+^; where ρ corresponds to the density (g/mL) and h corresponds to the levitation height (μm).

**Figure 2.**
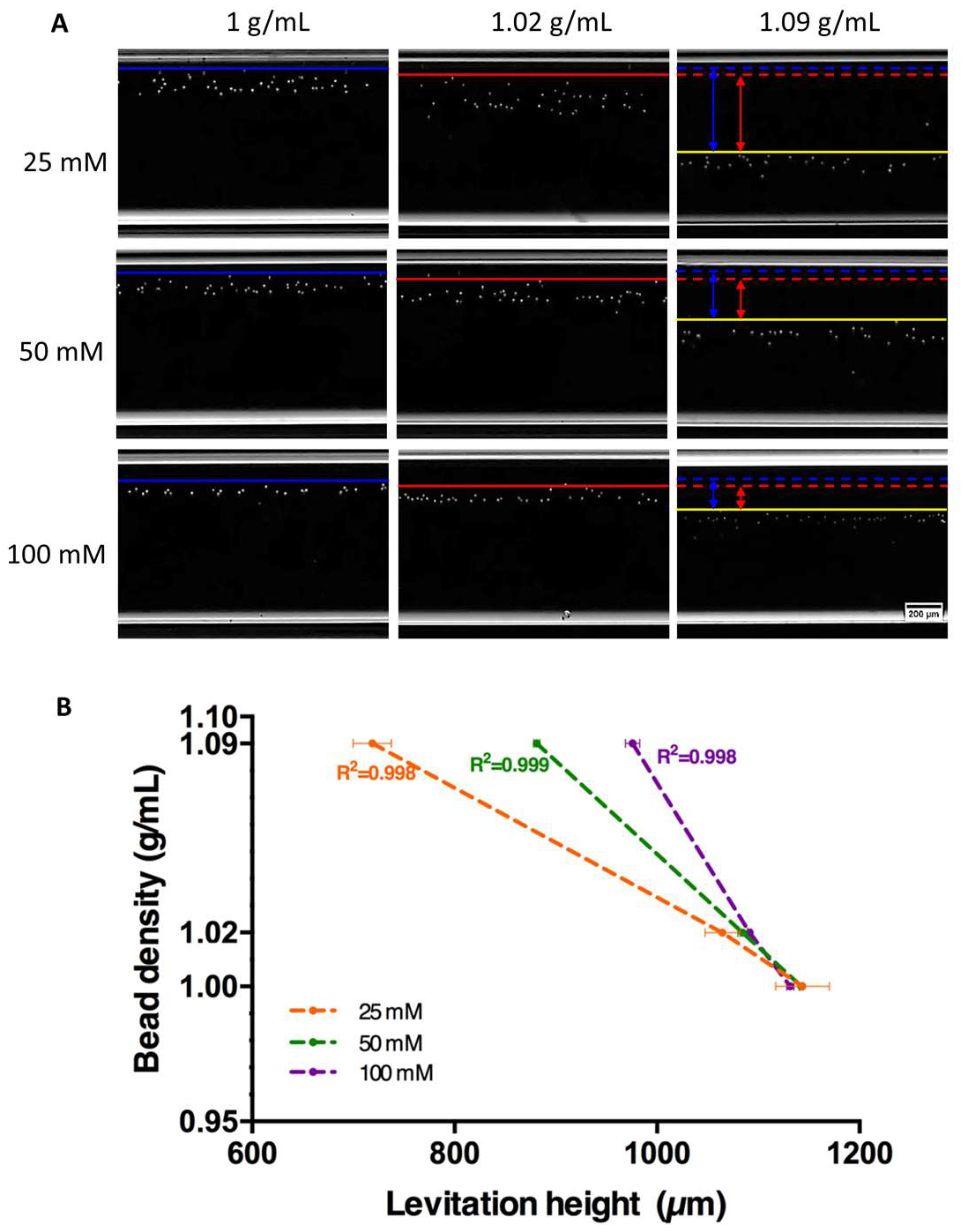
Calibration of the magnetic levitation system by using polymeric beads with known densities. (A) Images of the magnetically levitated beads with different densities (1 g/mL, 1.02 g/mL and 1.09 g/mL) in culture medium containing 25 mM, 50 mM or 100 mM Gd^3+^ solution. The blue line presents the uppermost level of the levitation heights of the beads with 1 g/mL density, the red line with 1.02 g/mL, and the yellow line with 1.09 g/mL. Blue and red arrows show the difference of levitation heights of the beads between density of 1.09 g/mL and, 1 and 1.02 g/mL. Scale bar: 200 μm. (B) Correlation between the levitation height and the density of beads. For each Gd^3+^ concentrations (25, 50 and 100 mM), a linear equation was obtained with determined levitation heights of beads (R^2^ =0.998, R2=0.999 and R2=0.998, respectively). Data is represented as mean ± SD.

### Detection of the adipogenic cells with microfluidic platform

D1 ORL UVA bone marrow mesenchymal stem cells were chosen as an initial experimental model for this study. D1 ORL UVA cells were first treated with an adipogenic induction medium for 15 days along with growth media controls (Supplementary Figure S2). Cells reached to confluency in both groups at the end of the first week. In contrast to the cells in growth medium, a fraction of cells treated with adipogenic induction started to accumulate lipid droplets at the end of the first week, and they continued to accumulate lipid droplets leading to hypertrophy. After 15 days of culture in induction medium, lipid accumulated cells were levitated in microfluidic device at 25 mM, 50 mM and 100 mM Gd^3+^ concentrations (Figure 3). In accordance with data obtained from polymeric beads, 100 mM Gd^3+^ resulted in a narrow band of 225 μm between average levitation height of high-density bulk cell population and levitation height of adipogenic cells with the lowest density. Decreasing the Gd^3+^ concentration increased this distance to 273 μm for 50 mM and 377 μm for 25 mM. Since the sensitivity of the levitation system to density changes in particles/cells increased as Gd^3+^ concentration decreased, we continued further experiments with 25 mM Gd^3+^ concentration. An in situ live / dead staining revealed that all cells, including cells that were large-sized and levitated at higher levels, appeared healthy (Supplementary Figure S3).

**Figure 3.**
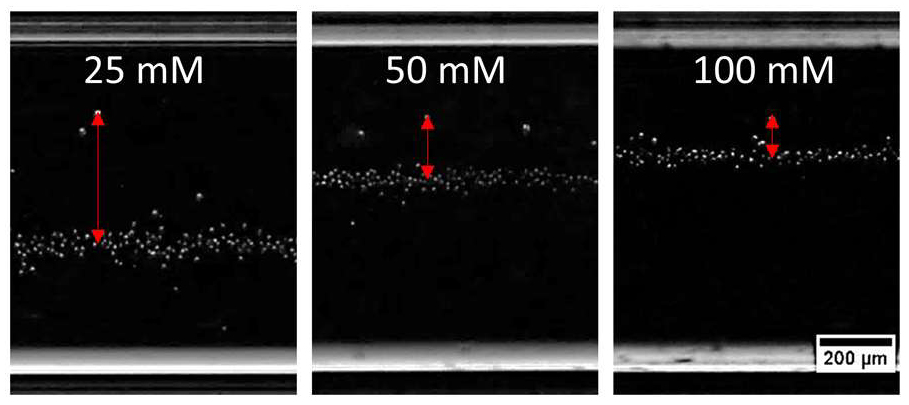
Optimization of paramagnetic medium concentration in the detection system. Magnetic levitation images of D1 ORL UVA cells cultured in adipogenic differentiation medium at the 15^th^ day using 25, 50 and 100 mM Gd^+3^ concentrations. Red arrows indicate height difference between average levitation height of general cell population and cells with lowest density, indicating the range of density based distribution. Scale bar: 200 μm.

### Determination of density profiles of adipogenic cells

In order to assess effect of adipogenic culture on levitation height, D1 ORL UVA cells were cultured with growth or adipogenic induction medium in plates up to 15 days. Cells then were levitated at 1^st^, 5^th^, 8^th^, 12^th^ and 15^th^ days of culture in the levitation system using 25 mM Gd^3+^ concentration, and changes in density of the cells depending on treatment time were analyzed (Figure 4). At the first day of culture, control and adipogenic cells resided in similar levitation heights, while at day 15 a fraction of cells subjected to adipogenic induction were positioned at higher levitation heights (Figure 4A). Cells in growth medium at day 1 were located at 761 ± 49 μm from the bottom magnet corresponding to density of 1.081 ± 0.010 g/mL according to the previously obtained equation for 25 mM Gd^3+^ concentration (Figure 4B). At day 15, control cells were similarly resided at 740 ± 31 μm corresponding to the density of 1.086 ± 0.005 g/mL. Cells in adipogenic induction medium had similar average density compared to control cells both at day 1 (p=0.96) and day 15 (p=0.33) (Figure 4C). Skewness of cell density distribution between groups were also similar between groups (p=0.94) at day 1 (Supplementary Figure S4A). We further assessed density in lower density fractions of cells, at day 1, average densities of cells in the lowest 5^th^ percentile for cells cultured in growth and adipogenic induction medium were similar also similar (p=0.71) (Supplementary Figure S4B). At day 15 however, average density of cells residing in the lowest 5^th^ percentile of density was 1.79% lower for adipogenic cells (p=0.03) than controls and 1.79% of the adipogenic population had density values smaller compared to the lowest density recorded (1.052 g/mL) on control cells (Supplementary Figure S4A). Furthermore, skewness of density distribution for cells cultured in growth and adipogenic medium were 0.99 ± 0.6 and -2.3 ± 0.6 (p=0.02), respectively (Supplementary Figure S4B). Physical changes in D1 ORL UVA cells during adipogenesis was not only limited to a reduction in density but also affected the cell size. Cells that had lower density than the average density of growth control cells (1.084 g/mL) at day 1 were analyzed for a measure of change in size during adipogenesis (Figure 5A-B). At the first day, there was no significant difference between quiescent and adipogenic induced cells that had 51 ± 18 μm^2^ and 36 ± 20 μm^2^ cell size in average, respectively (p=0.06). Cells in adipogenic induction had a slight but significant increase from 36 ± 19 μm^2^ at day 1 to 45 ± 28 μm^2^ at day 15 (p=0.05). Furthermore, there was a significant correlation between size and density of cells (R^2^=0.29) in contrast to undifferentiated cells (R^2^=0.01) after 15 days of culture (Figure 5C-D).

**Figure 4.**
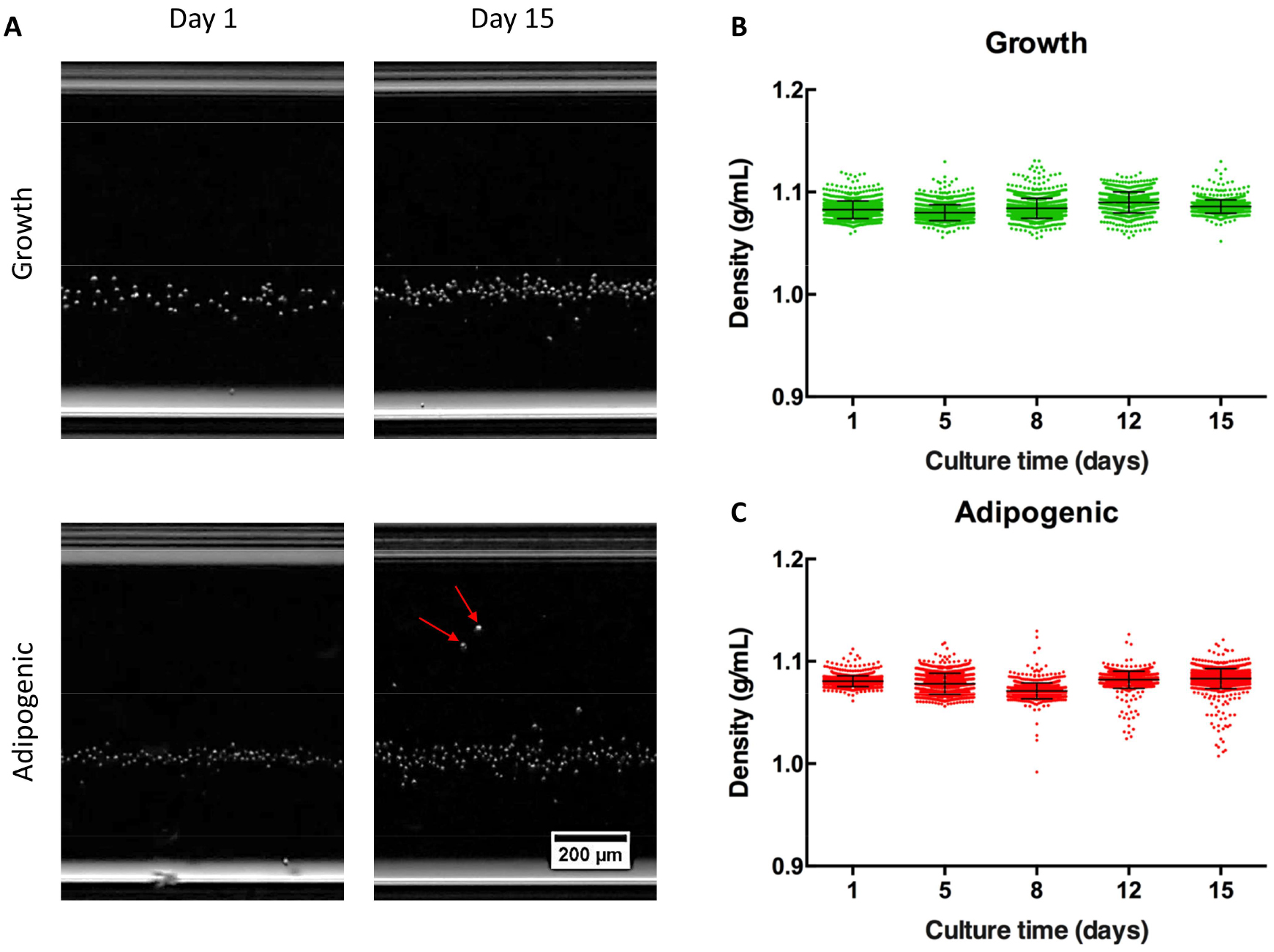
Levitation and density profiles of D1 ORL UVA cells using 25 mM Gd^3+^. (A) Levitation images of quiescent and adipogenic D1 ORL UVA cells at the 1^st^ and 15^th^ days. Arrows indicate differentiated cells with lower densities. Scale bar: 200 μm. (B, C) Scatter plots of densities of quiescent and adipogenic differentiated cells at the 1^st^, 5^th^, 8^th^, 12^nd^ and 15^th^ days. Data is represented as scattered with an inset of mean ± SD.

**Figure 5.**
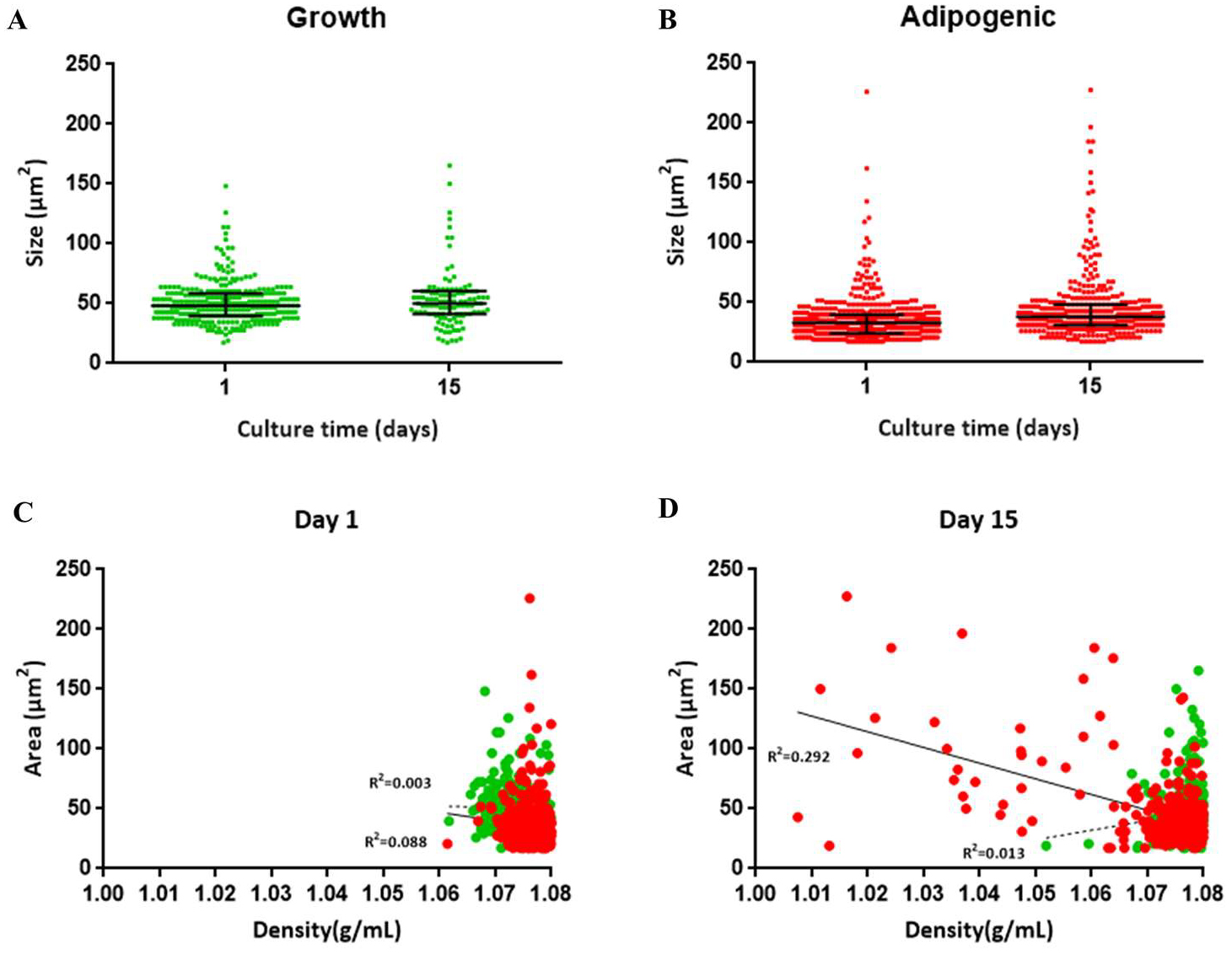
Alterations in cell size (area, μm^2^) during adipogenesis. (A, B) Size of quiescent and differentiated cells with density below 1.08 g/mL (average density of control cells) on day 1 and day 15 of culture. Data is represented as scattered with an inset of mean ± SD. (C, D) Relationship between size (area, μm^2^) and density (g/mL) of D1 ORL UVA cells cultured within growth and adipogenic medium for day 1 and day 15 of culture. Scatter plot represents relationship between density and size of differentiated and quiescent cells with density values < 1.08 g/mL. Dashed and solid lines indicate linear regression for quiescent and differentiated cells, respectively.

Since the lipid accumulation process of D1 ORL UVA cells were relatively slow and affected only a fraction of the population, we extended our study to adipogenesis in 7F2 cells, a bone marrow cell line that can accumulate high amount of lipids in relatively short time. Cells were cultured in similar growth medium or adipogenic differentiation medium for 10 days (Supplementary Figure S5), and levitated on 1^st^, 5^th^ and 10th days of culture with 25 mM Gd^3+^ (Figure 6). The cell density of the entire adipogenic cohort at 1^st^ day of culture was 1.079 ± 0.005 g/mL and decreased 1.76% (p<0.001) to 1.060 ± 0.029 g/mL after 10 days in adipogenic culture. Furthermore, average densities of cells in the 5^th^ percentile for 7F2 cells cultured in adipogenic induction medium for 10 days were observed as 1.003 ± 0.002 g/mL with minimum density of 0.989 g/mL and 37% of the adipogenic cell population had lower density compared to the lowest density cell observed in cells that were kept in the growth medium for 10 days.

**Figure 6.**
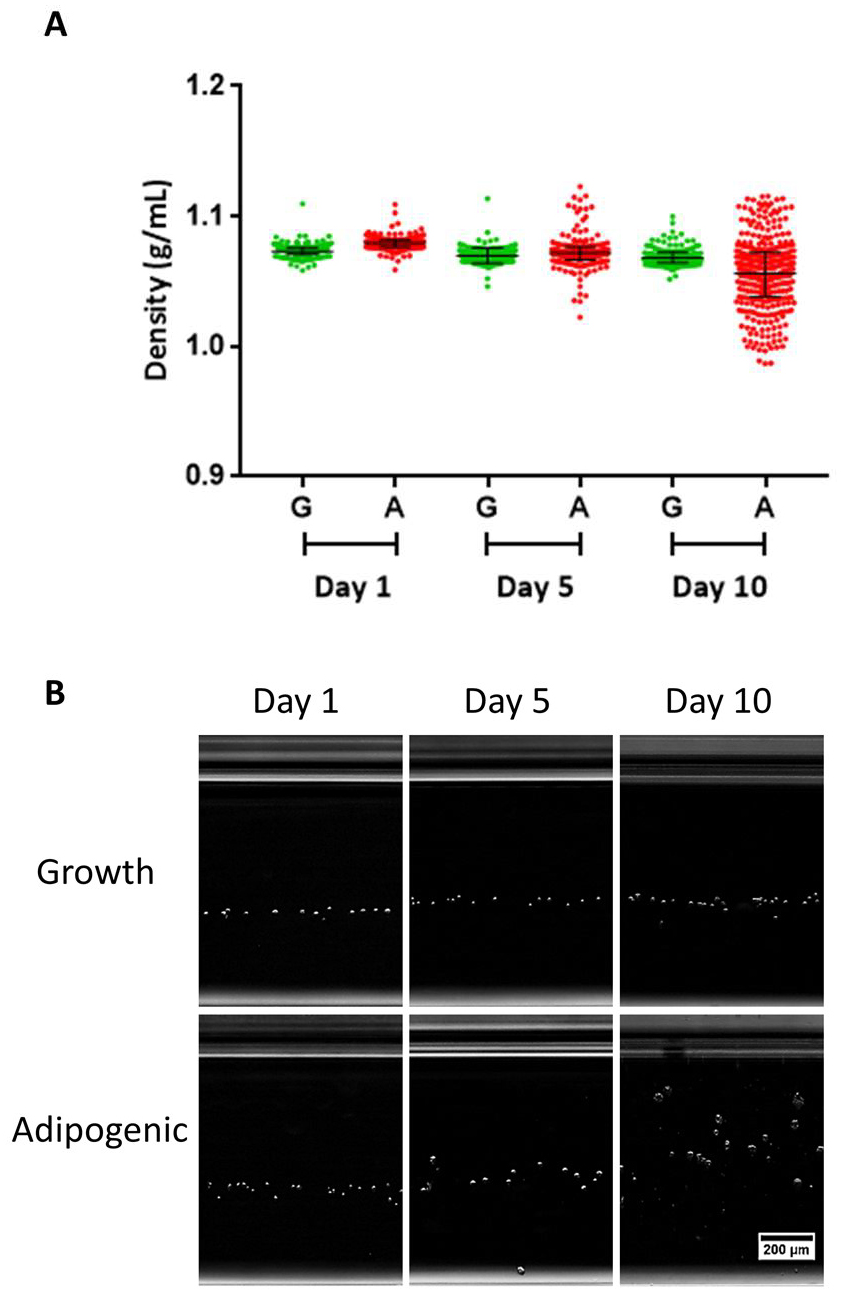
Levitation of 7F2 cells in magnetic levitation system. (A) Density dot plots of differentiated and undifferentiated cells at the 1^st^, 5^th^ and 10^th^ day of culture. Data is represented as scattered with an inset of mean ± SD. (B) Magnetic levitation (25 mM Gd^3+^) images of the cells cultured in the growth and adipogenic culture medium over 10 days. Scale bar: 200 μm.

### Detection of adipogenic differentiated cells mixed with stem cell population

In order to test whether the magnetic levitation system can be used in the detection of adipogenic cells in a mixed cell population, quiescent D1 ORL UVA cells (tracked with green fluorescent dye) and adipogenic-differentiated 7F2 cells (tracked with red fluorescent dye) were mixed at 1%, 5%, 10%, 25%, 50% (percentage of adipogenic 7F2 in the mix) ratios, and levitated with 25 mM Gd^3+^ containing medium (Figure 7A, Supplementary Figure S6). Results showed that increasing culture period resulted in an enhanced difference between the relative density of differentiated (7F2) and quiescent (D1) groups (Figure 7B). For the group in which quiescent cells and adipogenic cells were mixed at 50%, the average relative density of the cells obtained from the 1^st^ day of culture was similar (p=0.45), while 25.7% (p<0.001) difference was recorded between two cell groups at day 10. Similar trends were observed for other cell ratios. Even when the proportion of adipogenic cells in the quiescent cell population was reduced to 1%, low density 7F2 cells were distinctly observed in the detection system. To show that the differences observed in levitation height were based on differences in densities and not the cell type we levitated undifferentiated D1 ORL UVA and undifferentiated 7F2 cells (mixing ratio: 50%) at the 1^st^, 5^th^ and 10th day of culture as a control group (Supplementary Figure S7), showing the cells had similar density values (p>0.05) at all time points. Manual counting of 7F2 cells (red) in the heterogeneous mix resulted in determination of observed adipogenic cell fractions in different mixes. Compared to expected 7F2 fractions, observed values were mostly similar (p>0.05) as determined by chi-square test (Figure 7C). This similarity was more prominent in 7F2 cells that originate from 5^th^ and 10th days of adipogenic culture, where cells were committed to adipogenesis (Supplementary Figure S5).

**Figure 7.**
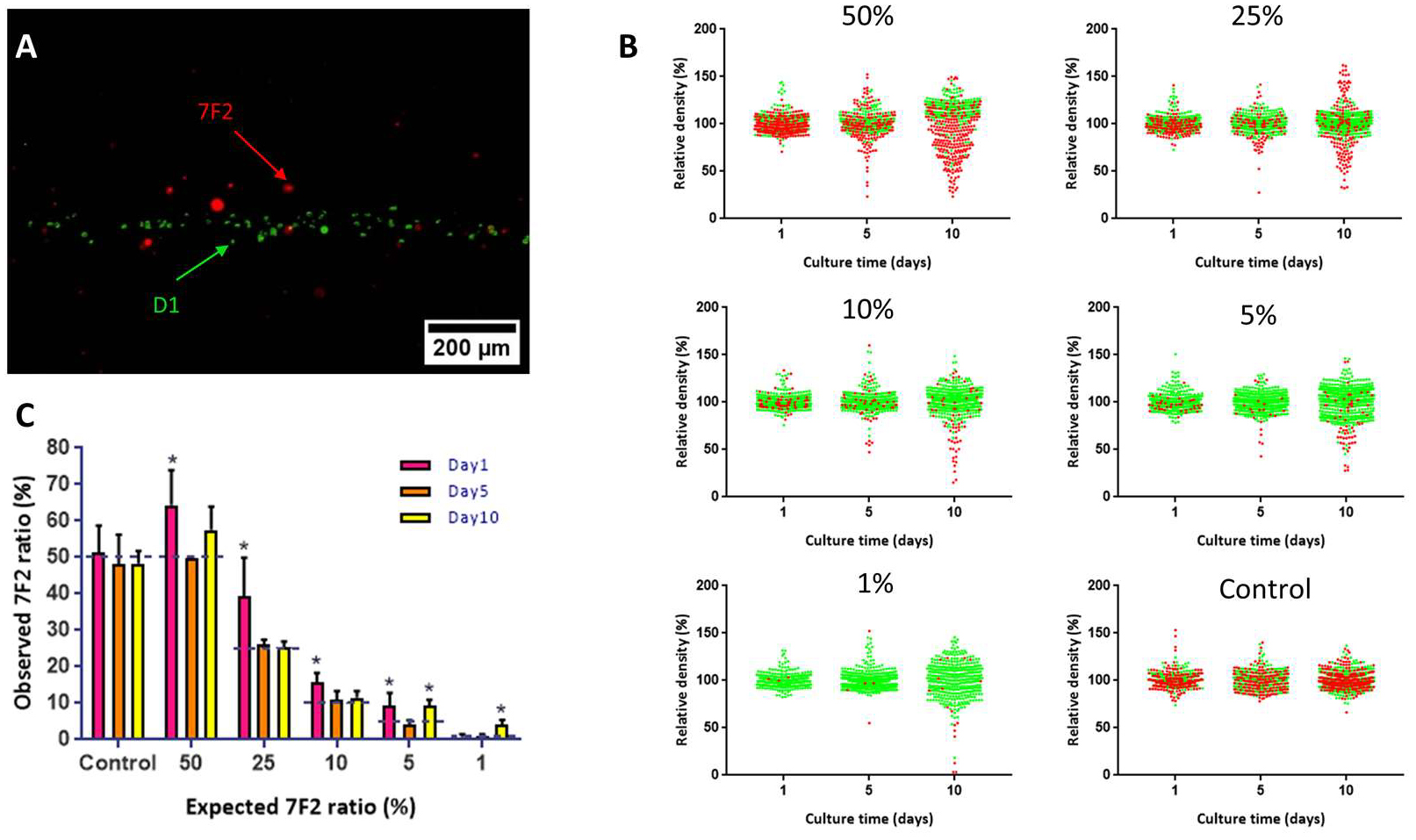
Detection of adipogenic cells in a heterogeneous cell population (A) Representative image of levitated quiescent D1 cells (green) and adipogenic differentiated 7F2 cells (red). Scale bar: 200 μm. (B) Scatter plot for relative density (%) of adipogenic differentiated 7F2 cells and quiescent D1 ORL UVA cells, cultured for 1, 5 and 10 days and mixed at different ratios of 7F2 cells (50%, 25%, 10%, 5% and 1%) and, undifferentiated 7F2 cells (red) and quiescent D1 ORL UVA cells with percentage of 50%, as control, for magnetic levitation (25 mM Gd^3+^). (C) Observed ratios of adipogenic differentiated 7F2 cells in these heterogeneous populations with magnetic levitation based detection system. Control group indicates undifferentiated 7F2 cells mixed with quiescent D1 ORL UVA cells with ratio of 50%. The chi-square test was performed for statistical analysis. Scale bar: 200 μm. Statistical significance was defined as *: p < 0.05; **: p<0.01; ***: p<0.001.

## DISCUSSION

In this study, we tested a novel method to detect the adipogenic differentiation of bone marrow stem cells based on cell density by magnetic levitation device. In summary, we firstly calibrated the detection system by using polymer beads with different densities to determine the correlation between levitation heights of particles and their density values depending on the concentration of the gadolinium contrast agent. Linear equations provided by the relationship between levitation heights and density values were used to measure single cell density. Secondly, we aimed to determine the optimum gadolinium concentration that should be used to perform density-based detection with D1 ORL UVA bone marrow stem cells, as a model. Eventually, 25 mM concentration was chosen for following experiments due to its high ability to distinguish adipogenesis-induced cells. Thirdly, the adipogenic cells in the stem cell population were successfully distinguished using the selected concentration in a heterogeneous population by magnetic levitation technology.

Magnetic levitation system has been previously reported as a detection/characterization method for cancer cells in a population containing different blood cell types, and as a drug testing system on prokaryotic cells ^60^. In this system, blood cells (i.e., red and white blood cells) were located at lower levitation heights with higher density and different type of cancer cells such as breast adenocarcinoma (MDA-MB-231), esophageal adenocarcinoma (JHEsoAD1), colorectal adenocarcinoma (HT29) were positioned at higher levels as a result of their lower density values. On the other hand, the effect of the antibiotic treatment, known to cause cellular composition changes of bacterial cells, on the density of cells was shown in real time by the levitation system ^60, 68^. Here, we aimed to adapt the system to detect adipocytes by using separation function of the principle depending on the density. In conclusion, the levitation system was successfully performed for the detection of adipogenic differentiated stem cells. The density of D1 ORL UVA stem cells and 7F2 osteoblast cells (>1.07 g/mL) is strongly consistent with the density of previously reported bone marrow-derived stem cells ^69, 70^. On the other hand, adipogenic cells were found to be less dense (down to 0.989 g/mL) than all these cell types, regardless of the origin of adipocytes. These results show that the magnetic levitation system has a potential to distinguish different cell populations in the bone marrow. Even considering the suitability of this system for cell culture ^64^, it is also promising that these cell types can be levitated and cultured at different levels in a single culture chamber and the interactions between these cells can be examined at the cellular and molecular level.

The density range of the adipose tissue is determined as 0.925 and 0.970 g/mL ^71^, however density values of adipocytes at single cell level are not yet available in the literature. However, some density measurement techniques for single cells can be used to measure density of adipocytes, such as OEK platform, SMR. Contrary to these methods that measure single cell density by the help of cell mass and volume analyzed with complex and expensive instrumentations, our system offers a fast and cost-effective way to measure densities by using a specific equation via magnetic levitation system ^57, 58, 72^. Besides, most of the microfluidic systems for density measurement such as SMR, multiphase stems, usually need two or more liquids with different density ^49, 54, 57^. Magnetic levitation technique does not need multiple phases with different liquids and thus offer ease of application. In addition, unlike other methods that allow analysis at the single cell level (i.e. density gradient centrifugation), magnetic levitation platform provides density measurement of cells in great quantities at the same time ^59^. Another advantage of this principle is that it allows label-free detection and it is possible to adapt the system to separate cells without modifications on them for following studies.

We achieved measuring density of adipogenic differentiated cells and findings revealed that lipid accumulated cells had positioned at higher levels than undifferentiated cells at single cell level. It was thought that differences in levitation heights of adipocytes could be caused by either differences in state (lineage) or the amount of accumulated lipid droplets of differentiated cells. Thereby, our detection system has a potential to become an informative method related to lipid accumulation and cell state in contrast to current methods such as optical microscopy. Additionally, determination of adipocyte size is important for adipogenesis and metabolic studies since the increase in cell size of adipocytes is one of the parameters of adipogenesis and also adipocyte size influences cellular metabolism rate ^32, 33, 73^. Some metabolic functions such as secretion of cytokines by adipocytes are believed to be related with adipocyte size and size changes is relatable to disorders such as obesity, insulin resistance and type 2 diabetes ^74, 75^. Previous techniques for determination of the size of adipocytes are generally time-consuming and terminal that relies on cell fixation and then monitoring cell images obtained with a camera ^73, 76^. In contrast to these systems, our system provide simultaneous monitoring of relationship between cell density and cell size as evidenced by the inverse correlation between size and density in adipocytes during the lipid accumulation process.

Besides the targeted cellular changes to be tested (i.e. differentiation), we also noticed that the culture conditions may affect the density of the cells. In this study, we tried longer term culturing of D1 ORL UVA cells to 3 weeks to increase adipogenic fraction. But this resulted in non-homogenous distribution of both control and adipogenic cells in levitation system, potentially from reduced cellular health and confined cell sizes (data not shown). Another issue to be considered in the determination of density by magnetic levitation system is the magnetic susceptibility of the paramagnetic medium. Although 25 mM Gd^3+^ is appropriate to distinguish the cell types, stem cells and adipocytes used here, this value may be required to be customized for high resolution detection in cell combinations with different range of density.

In conclusion, our method provides a label-free, real-time detection system for adipogenic differentiation based on their density. A large number of adipocytes can be detected, and their density and size can be measured at single cell level simultaneously. This protocol allows a fast and easy way to detect mature adipocytes with low density that is not possible with other methodologies. The density-based protocol established here may offer a wide range of applications including drug discovery and tissue engineering. Although the present system is limited to detection, the platform has a potential to be modified into a separation system based on density by providing flow of the sample to isolate cell groups levitated at distinct heights for further culture or downstream analysis.

## MATERIALS AND METHODS

### Experimental Setup

A magnetic levitation device composed of micro-capillary channel (1-mm × 1-mm cross-section, 50-mm length, and 0.2-mm wall thickness, Vitrocom) between two N52-grade neodymium magnets (NdFeB) (50-mm length, 5-mm height and 2-mm width, Supermagnete), was constructed to create a magnetic field gradient perpendicular to gravity and thus levitate cells in the paramagnetic medium and two parallel mirrors (Thorlabs) were placed at 45° to visualize levitation of cells using an inverted microscope (Olympus IX-83) ^64^.

### Cell Culture

D1 ORL UVA (mouse bone marrow stem cells) and 7F2 (mouse osteoblasts) were obtained from ATCC. Cells were grown in a growth medium supplemented with 10% fetal bovine serum (FBS) and 1% penicillin/streptomycin at 5% CO_2_ humidified atmosphere at 37°C. D1 ORL UVA and 7F2 cells were cultured in Dulbecco’s Modified Eagle’s medium (DMEM high glucose, Gibco) and alpha modified essential medium (αMEM), respectively. The growth medium was refreshed every 2-3 days and cells were passaged every 4-6 days. For differentiation of D1 ORL UVA cells into adipocytes, 1000 cells/well was seeded in 24-well plates and after 48h, adipogenesis was induced by differentiation medium that contains 10 mM dexamethasone, 50mM indomethacin, 5×10^-3^ mg/ml insulin for 15 days. Likewise, 7F2 cells were induced in αMEM with induction agents for 10 days. The adipogenic induction medium was replaced every 2-3 days. The cells were imaged at 10× under an inverted microscope (Olympus IX-83).

### Magnetic levitation of polymeric beads

Polymer beads with different densities; 1 g/mL, 1.02 g/mL (with size of 10-20 μm) and 1.09 g/mL (with size of 20-27 μm) (Cospheric LLC., ABD), were levitated in the culture medium containing 25 mM, 50 mM or 100 mM gadolinium (Gd^3+^) (Gadavist^®^, Bayer). Levitated beads were visualized at 4× under an inverted microscope (Olympus IX-83) after beads reached the equilibrium position (within ^~^10 min) in the magnetic levitation platform. The levitation heights of the beads (distance from the upper limit of the bottom magnet) were determined using the ImageJ Fiji software.

### Magnetic levitation of the cells

D1 ORL UVA cells that were cultured in growth and adipogenic induction medium were trypsinized at the 1^st^, 5^th^ 8^th^,12nd and 15^th^ days, and were centrifuged at 125 × g for 5 min. Pellet was resuspended to 10^5^ cells/ml in the culture medium containing Gd^3+^ paramagnetic agent with concentrations of 25, 50 and 100 mM. Then, 50 μL samples (5000 cell/capillary) were loaded into the microcapillary channel. The samples were levitated until the cells reached the equilibrium position (^~^15 min) and imaged under the microscope. Later on, levitation images were analyzed by ImageJ Fiji software to determine levitation heights/density and cell size. Similarly, 7F2 cells were trypsinized at the 1^st^, 5^th^ and 10th days and levitated using 25 mM Gd^3+^ concentration within the levitation system. Then, the same analysis method was applied for measuring levitation heights of cells and cell size.

### Live/Dead Assay

D1 ORL UVA cells were seeded at a concentration of 1 × 10^3^ cells/well into a 24-well plate and cultured for 22 days. Cell viability assay (calcein-AM/propidium iodide, Sigma Aldrich) was carried out to test the viability of levitated cells. The cells were stained for 15 min and levitated using three different concentrations of Gd^3+^ (25, 50 and 100 mM). Then they were imaged under the fluorescence microscope (Olympus IX-83).

### Fluorescent Staining and Mixing of D1 ORL UVA and 7F2 Cells with Different Ratios

D1 ORL UVA was cultured with quiescent medium and trypsinized at 1, 5 and 10th days. The cells were suspended at a density of 1 × 10^6^ cells/mL in serum-free DMEM culture medium. Then, they were stained with 5 μM of DiO (green) cell-labeling solution (Vybrant^TM^). After incubation at 37°C for 20 min, the labeled cell suspensions were centrifuged at 1500 rpm for 5 min and the supernatant was removed. The washing procedure was repeated two more times. Likewise, 7F2 cells were cultured in αMEM growth medium and adipogenic induction medium. After trypsinization, the cells were suspended at the same concentration in serum-free culture medium and stained with 5 μM of DiI (red) cell-labeling solution using the same protocol. Labeled cells were mixed at different percentages of 50%, 25%, 10%, 5% and 1%, levitated in magnetic levitation system and imaged under the fluorescence microscope (Olympus IX-83).

### Statistical analysis

During the study, all experiments were repeated at least three times. Data on density and levitation height were presented with the mean and standard deviation (mean ± SD) or with the scatter plots and median with interquartile range. Student’s t-test (two-tail) or one-way ANOVA with Tukey’s multiple comparisons test were used to determine statistical significance and P < 0.05 was considered to be statistically significant. The Chi-square test was used to test the associations between observed and expected cell ratios in magnetic levitation device. Graphs showing levitation height versus density of beads at different Gd^3+^ concentrations were plotted and linear regression over the data was performed to obtain equations providing density of levitated particles/cells levitated in magnetic detection system.

## ACKNOWLEDGMENTS

Financial support from The Scientific and Technological Research Council of Turkey (215S862 - EO, 116M298 - HCT) and Turkish Academy of Sciences (Young Investigator Award - EO) is gratefully acknowledged. We thank Sena Yaman for fabricating the magnetic levitation platform. We are also thankful for the helpful discussions with Ozden Yalcın-Ozuysal, PhD.

## AUTHOR CONTRIBUTIONS

E.O., H.C.T. and G.M. conceived and designed the study; O.S. and M.A. performed the experiments; O.S. and M.A. and E.O. analyzed the data; O.S., M.A. G.M., H.C.T. and E.O. wrote the manuscript.

## COMPETING INTERESTS

The authors declare no competing interests.

## SUPPLEMENTARY FIGURES

**Supplementary Figure S1.**
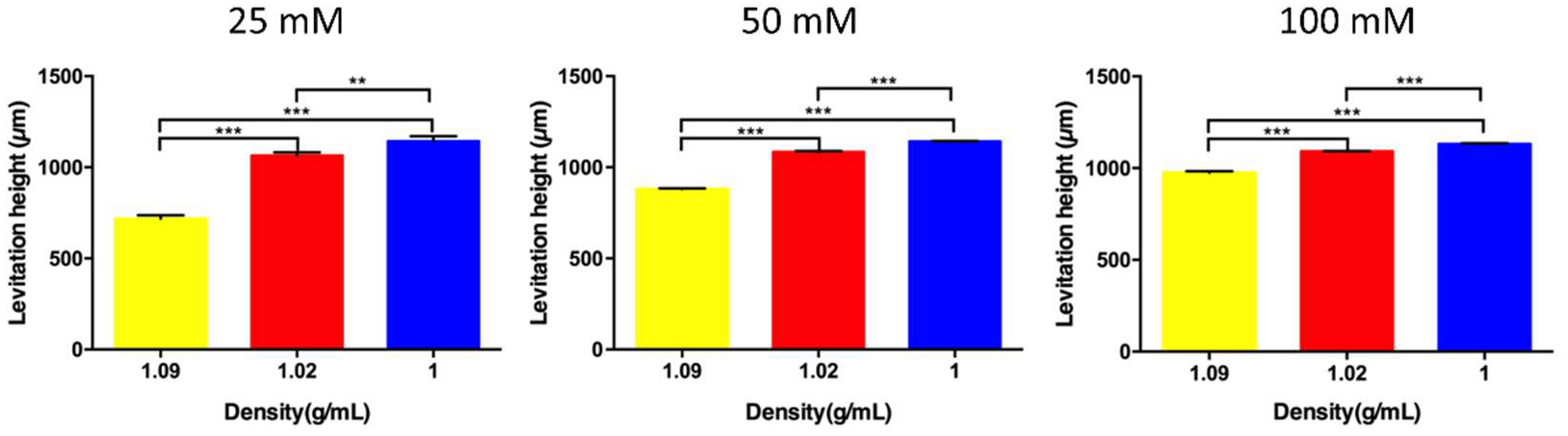
Levitation heights of beads with different densities (1, 1.02 and 1.09 g/mL) at 25, 50 and 100 mM Gd^3+^ containing medium. Data are presented as mean of replicates with error bars (±SD) and one-way ANOVA with Tukey’s multiple comparisons test was performed for statistical analysis. Statistical significance was defined as *: p < 0.05; **: p<0.01; ***: p<0.001.

**Supplementary Figure S2.**
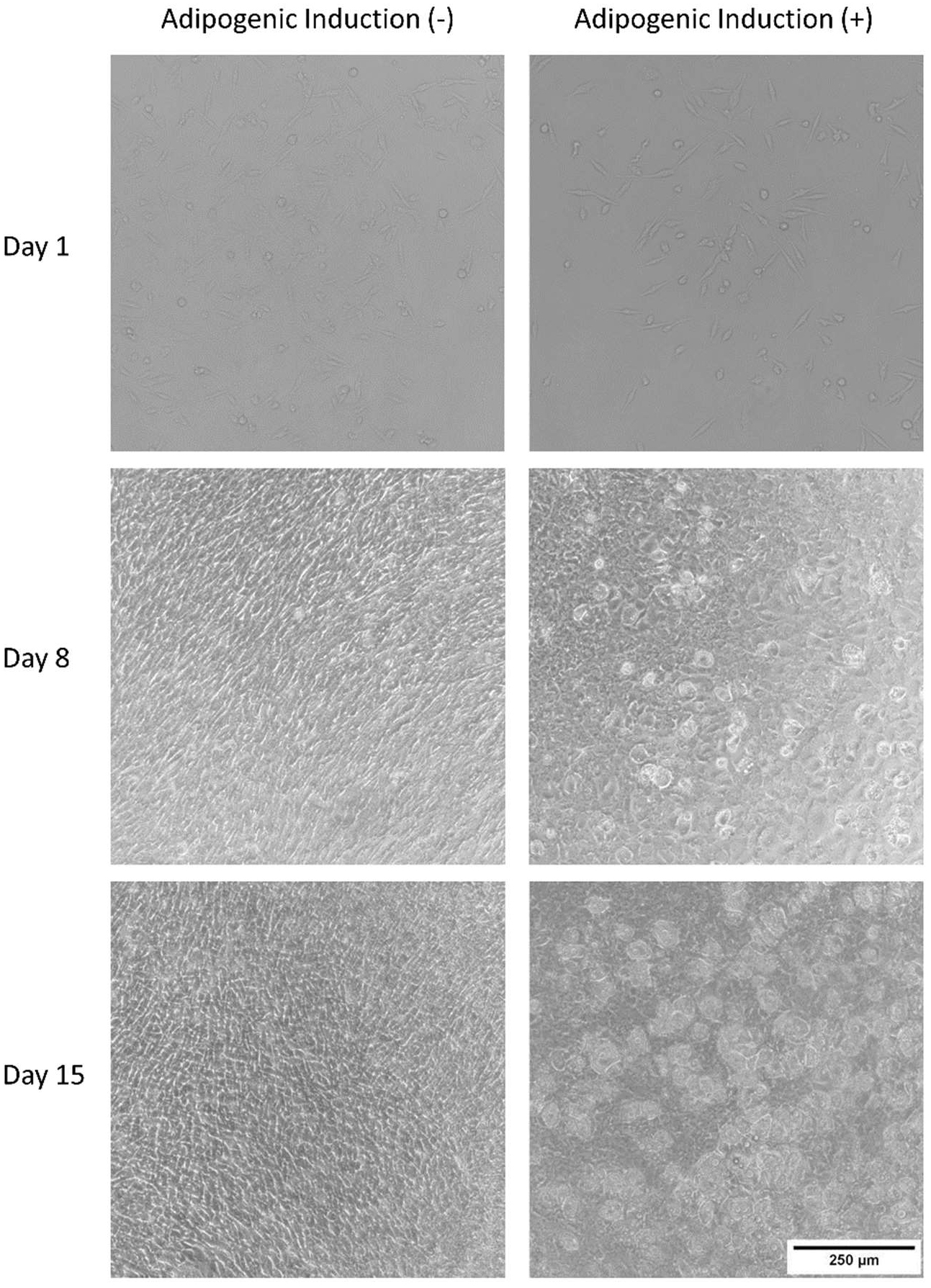
Cell culture images of D1 ORL UVA cells that were cultured in standard growth medium and adipogenic differentiation medium over 15 days. Scale bar: 250 μm.

**Supplementary Figure S3.**
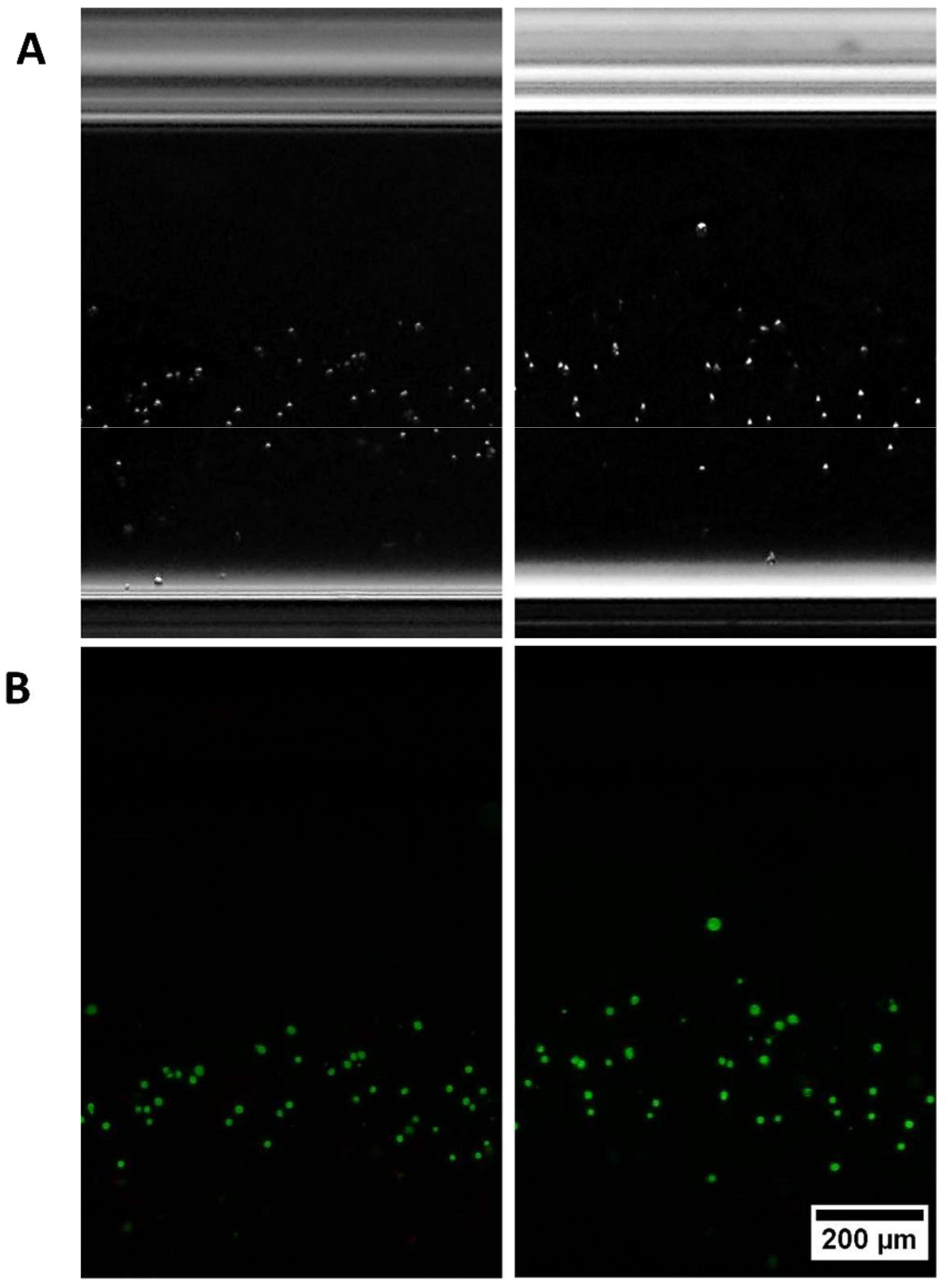
Bright-field (A) and fluorescent (B) microscopy images of D1 ORL UVA cells cultured in growth or adipogenic differentiation medium at the 22^nd^ day, in magnetic levitation device (25 mM Gd^3+^). Cell viability was visualized by live-dead (live: green, dead: red) staining. Dead cells were not detected in the system. Scale bar: 200 μm.

**Supplementary Figure S4.**
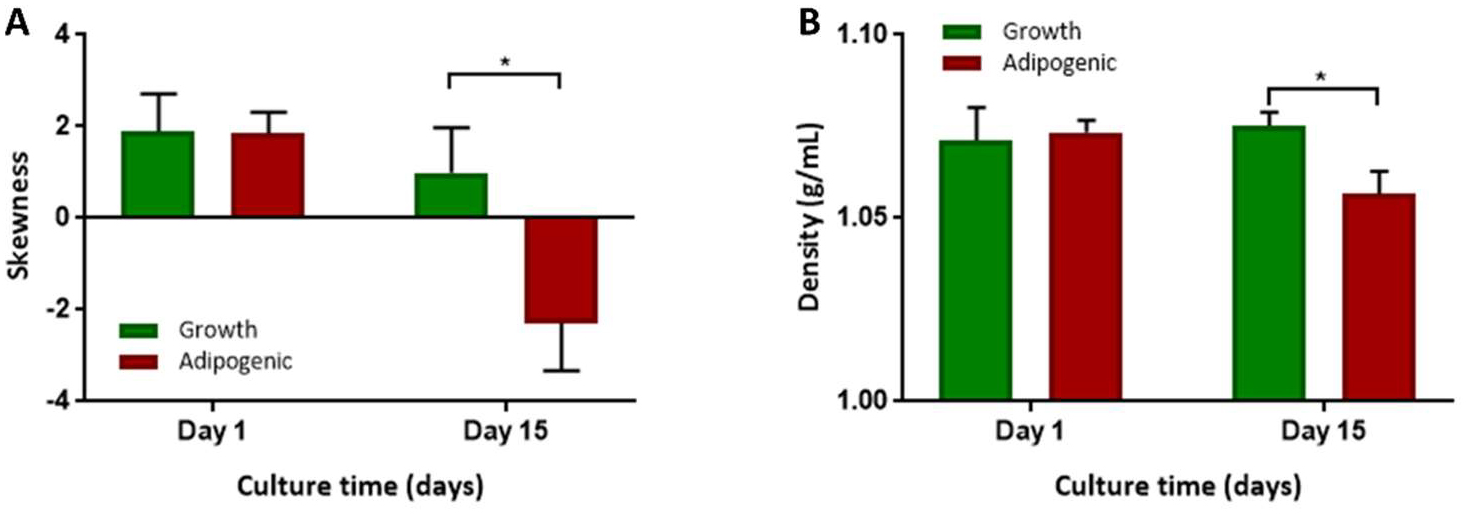
(A) Skewness of density values of control and adipogenic populations at the 1^st^ and 15^th^ days. (B) Average density values of lowest 5^th^ percentile pertaining to control and adipogenic populations. Data is represented as mean ± SD. *: p < 0.05; **: p<0.01; ***: p<0.001.

**Supplementary Figure S5.**
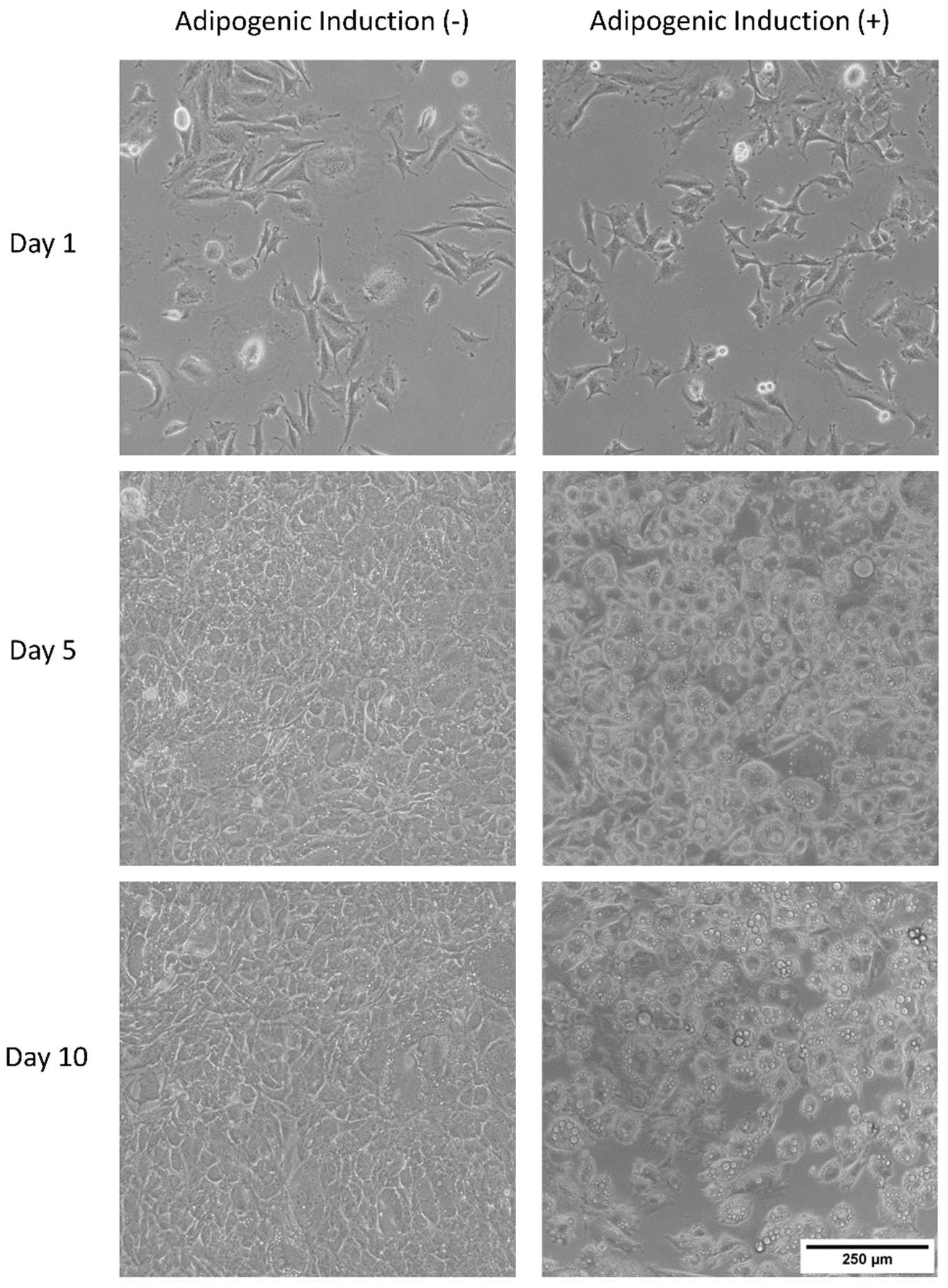
Cell culture images of 7F2 cells that were cultured in standard growth medium and adipogenic medium for 10 days. Scale bar: 250 μm.

**Supplementary Figure S6.**
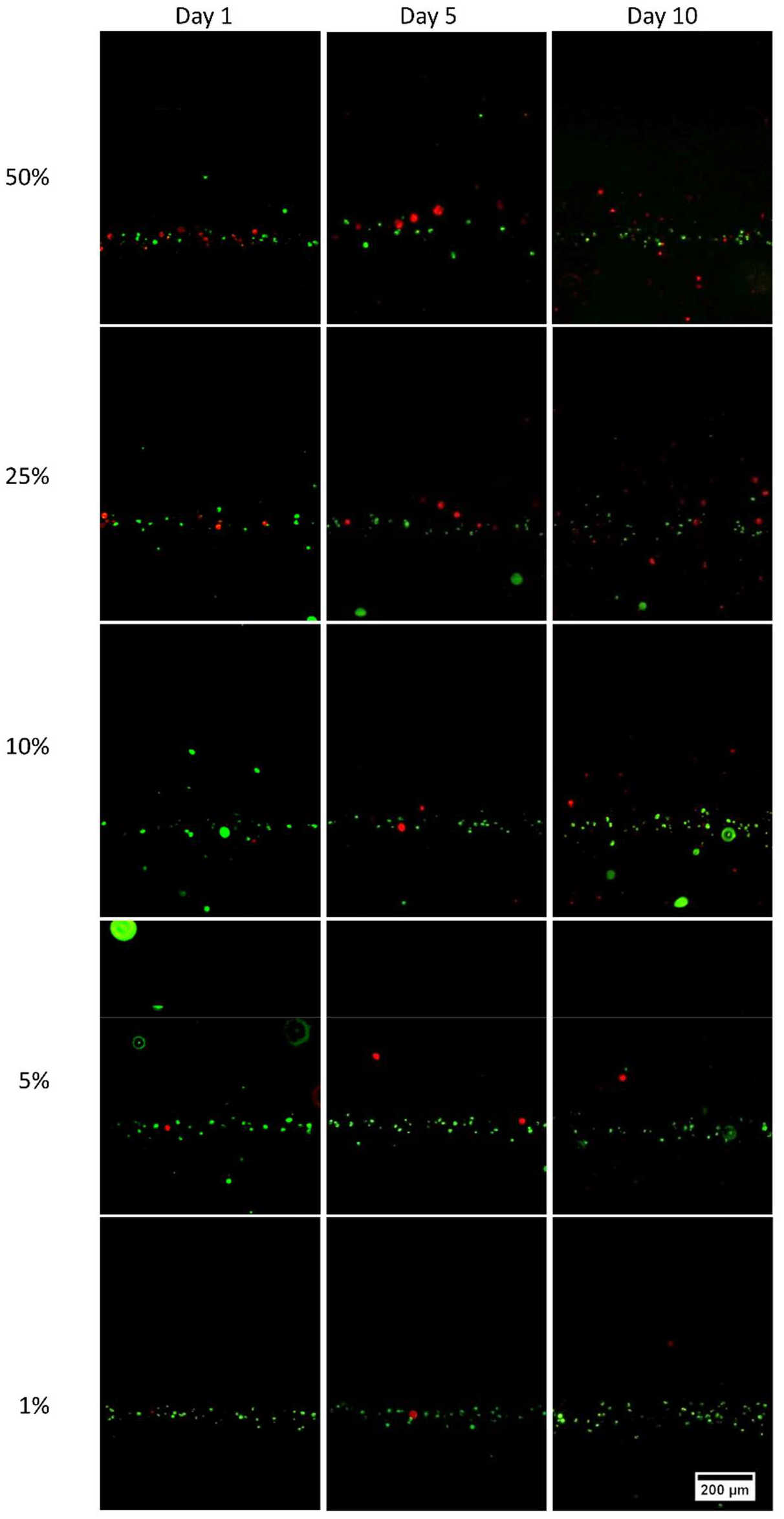
Fluorescent images of the levitated heterogeneous populations consisted of quiescent D1 ORL UVA cells (green) and adipogenic differentiated 7F2 cells (red) at different ratios (50%, 25%, 10%, 5% and 1%) after 1, 5 and 10-day culture. Scale bar: 200 μm.

**Supplementary Figure S7.**
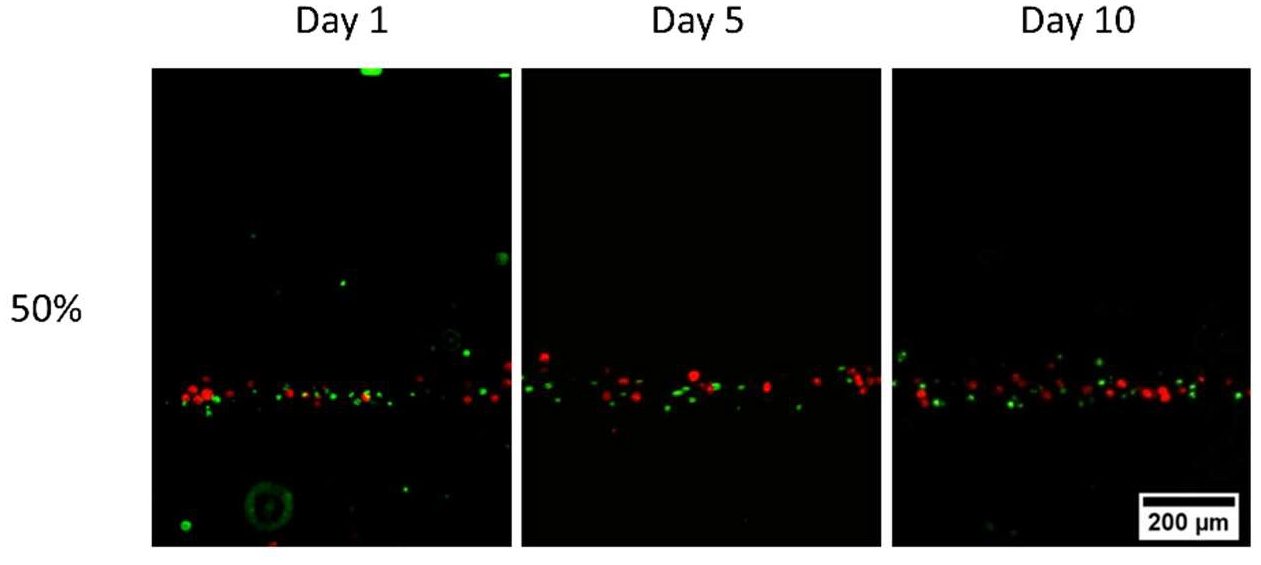
Fluorescent images of the levitated heterogeneous population consist of undifferentiated 7F2 (red) cells and D1 ORL UVA (green) cells mixed at 50% ratio after 1, 5 and 10-day culture. Scale bar: 200 μm.

